# Embryonic origin and serial homology of gill arches and paired fins in the skate (*Leucoraja erinacea*)

**DOI:** 10.1101/2020.07.02.183665

**Authors:** Victoria A. Sleight, J. Andrew Gillis

## Abstract

Paired fins are a defining feature of the jawed vertebrate body plan, but their evolutionary origin remains unresolved. Gegenbaur proposed that paired fins evolved as gill arch serial homologues, but this hypothesis is now widely discounted, owing largely to the presumed distinct embryonic origins of these structures from mesoderm and neural crest, respectively. Here, we use cell lineage tracing to test the embryonic origin of the pharyngeal and paired fin skeleton in the skate (*Leucoraja erinacea*). We find that while the jaw and hyoid arch skeleton derive from neural crest, and the pectoral fin skeleton from mesoderm, the gill arches are of dual origin, receiving contributions from both germ layers. We propose that gill arches and paired fins are serially homologous as derivatives of a continuous, dual-origin mesenchyme with common skeletogenic competence, and that this serial homology accounts for their parallel anatomical organization and shared responses to axial patterning signals.

## Introduction

It was classically proposed that the paired fins of jawed vertebrates evolved by transformation of a gill arch – a theory stemming largely from Gegenbaur’s (1878) interpretation of a shared anatomical ground plan between the gill arch and pectoral fin skeletons of cartilaginous fishes (sharks, skates and rays) (reviewed by Coates, 1994; 2003). In vertebrate embryos, the jaw, hyoid and gill arch skeleton (or, in amniotes, their derivatives, the jaw, auditory ossicles and laryngeal skeleton) arises from a series of transient, bilaterally paired pharyngeal arches that form on the sides of the embryonic head (Gillis *et al*., 2012; Graham *et al*., 2019), while the paired fins or limbs of jawed vertebrates arise as buds that project from the embryonic trunk (Tickle, 2015). Cell lineage tracing studies in bony vertebrates (Chai *et al*., 2000; Jiang *et al*., 2002; Couly *et al*., 1993; Kague *et al*., 2012) have revealed that the pharyngeal arch skeleton derives largely from the neural crest – a vertebrate-specific, multipotent cell population that undergoes epithelial-to-mesenchymal transition from the dorsal neural tube, and that gives rise to a plethora of derivatives, including skeletal and connective tissue lineages (Green *et al*., 2015). The skeleton of paired appendages, on the other hand, derives from the lateral plate – a distinct mesodermal subpopulation that arose along the chordate stem (Prummel *et al*., 2019). As shared embryonic origin has classically been regarded as a key criterion for serial homology (discussed by Hall, 1995), Gegenbaur’s gill arch hypothesis of paired fin origin was widely discounted, and is now generally deemed fundamentally flawed (Coates, 2003).

Importantly, though, the distinct embryonic origins of the gill arch and paired fin skeletons may not hold true: mesodermal contributions to the posterior pharyngeal skeleton have been demonstrated in tetrapods, but are much less widely appreciated than those from neural crest. Cell lineage tracing using quail-chick chimaeras has revealed that the avian cricoid and arytenoid laryngeal cartilages derive from lateral mesoderm, and not neural crest (Noden, 1986, 1988; Evans and Noden, 2006) – a finding that has since been corroborated by genetic lineage tracing experiments in mouse (Tabler *et al*., 2017; Adachi *et al*., 2020). Additionally, ablation (Stone, 1926) and lineage tracing experiments (Davidian and Malashichev, 2013; Sefton *et al*., 2015) have revealed a mesodermal origin of the posterior basibranchial cartilage in axolotl. Currently, however, there are no mesodermal fate maps of the pharyngeal skeleton of fishes, and so it remains to be determined whether mesodermal contributions to the posterior pharyngeal skeleton are an ancestral feature of jawed vertebrates, and whether mesoderm is competent to give rise to gill arch cartilages – i.e. the ancestral skeletal derivatives of the posterior pharyngeal arches, and Gegenbaur’s proposed evolutionary antecedent to paired fins.

We sought to map the contributions of neural crest and mesoderm to the pharyngeal and paired fin endoskeleton in a cartilaginous fish, the little skate (*Leucoraja erinacea*), as data from this lineage will allow us to infer ancestral germ layer contributions to the pharyngeal and paired fin skeletons, and to test the developmental potential of neural crest and mesodermal skeletal progenitors in a taxon that has retained an ancestral gill arch anatomical condition. We find that the gill arch skeleton of skate embryos receives contributions from both cranial neural crest and lateral mesoderm, revealing its dual embryonic origin. These findings point to an ancestral dual embryonic origin of the pharyngeal endoskeleton of jawed vertebrates, and to gill arches and paired appendages as serial derivatives of a dual-origin, neural crest- and mesodermally-derived mesenchyme with equivalent skeletogenic potential at the head-trunk interface.

## Results

### Neural crest and lateral mesoderm in the skate neurula

In the skate, neural tube closure begins at embryonic stage (S)16 and is complete by S18 (Ballard *et al*., 1993). *in situ* expression analysis of the gene encoding the conserved neural crest specifier Foxd3 reveals that by S18, pre-migratory cranial neural crest cells are specified within the dorsal neural tube but are not yet undergoing epithelial-to-mesenchymal transition (Fig. 1A). At S18, we can also recognize molecularly distinct lateral mesodermal populations, including *tbx1*-positive cranial paraxial mesoderm (Fig. 1B), which is morphologically continuous with *pitx2*- and *hand2*-positive lateral plate mesoderm (Fig. 1C), and *myf5*-positive somitic and pre-somitic paraxial mesoderm (Fig. S1). The clear spatial segregation and accessibility of these tissues in S18 skate embryos (Fig. 1D) renders them amenable to fate mapping by labelling with lipophilic dyes – either by microinjecting the lumen of the neural tube (to label pre-migratory neural crest cells), or by microinjecting mesodermal mesenchyme underneath the head ectoderm – and so we used this approach to directly test the contributions of these tissues to the pharyngeal and paired fin endoskeleton.

**Figure 1:**
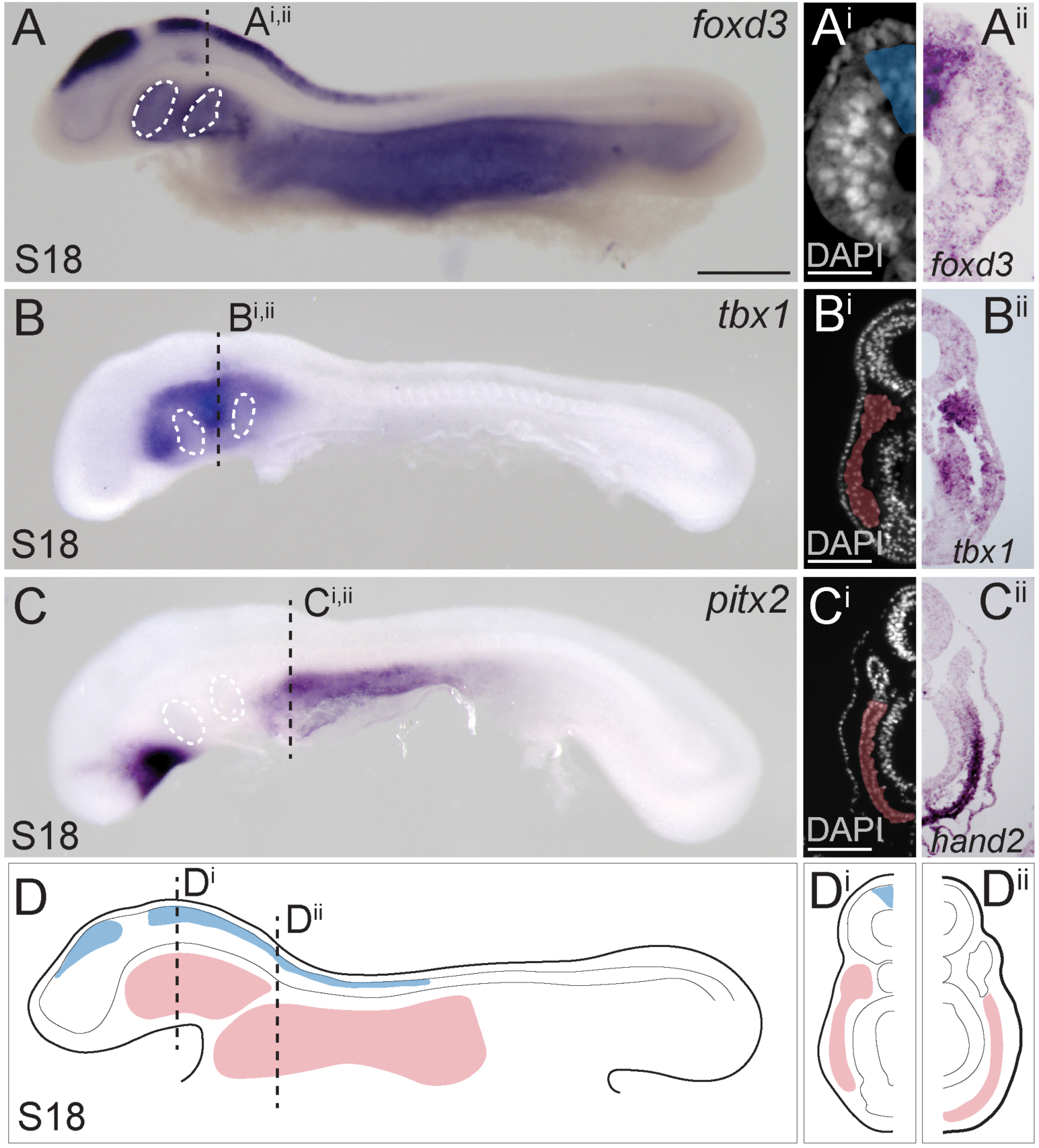
Neural crest and mesoderm after skate neurulation. **A)** Wholemount mRNA *in situ* hybridization for *foxD3* reveals expression in **A**^**i**^, **A**^**ii**^) pre-migratory neural crest cells within the dorsal neural tube of the skate embryo at S18. **B, B**^**i**^, **B**^**ii**^) *tbx1*-expressing head mesoderm grades into **C)** *pitx2-* and **C**^**i**^, **C**^**ii**^) *hand2*-expressing lateral plate mesoderm in the skate embryo at S18. **D)** Schematic representation of neural crest, head mesoderm and lateral plate mesoderm tissues targeted for cell lineage tracing in this study. Scale bars: **A, B, C** = 700μm; **A**^**i**^ = 65μm; **B**^**i**^, **C**^**i**^ = 120μm.

### Neural crest contributes to the skate jaw, hyoid and gill arch skeleton

To label skate cranial neural crest (NC) cells, we microinjected the lumen of the neural tube with CM-DiI at the hindbrain level. This resulted in very bright labelling at the point and time of injection (Fig. 2A), though analysis of embryos collected shortly post-injection in section reveals that the cells of the neural tube were labelled around its entire circumference, broadly, along the length of the hindbrain region (Fig. 2A^i^). At five days post-injection, we observed abundant CM-DiI-labelled NC cells streaming into the pharyngeal arches (Fig. 2B), and at S31/32 (∼8-10 weeks post-injection), we tested for NC contributions to cartilages throughout the pharyngeal skeleton.

**Figure 2:**
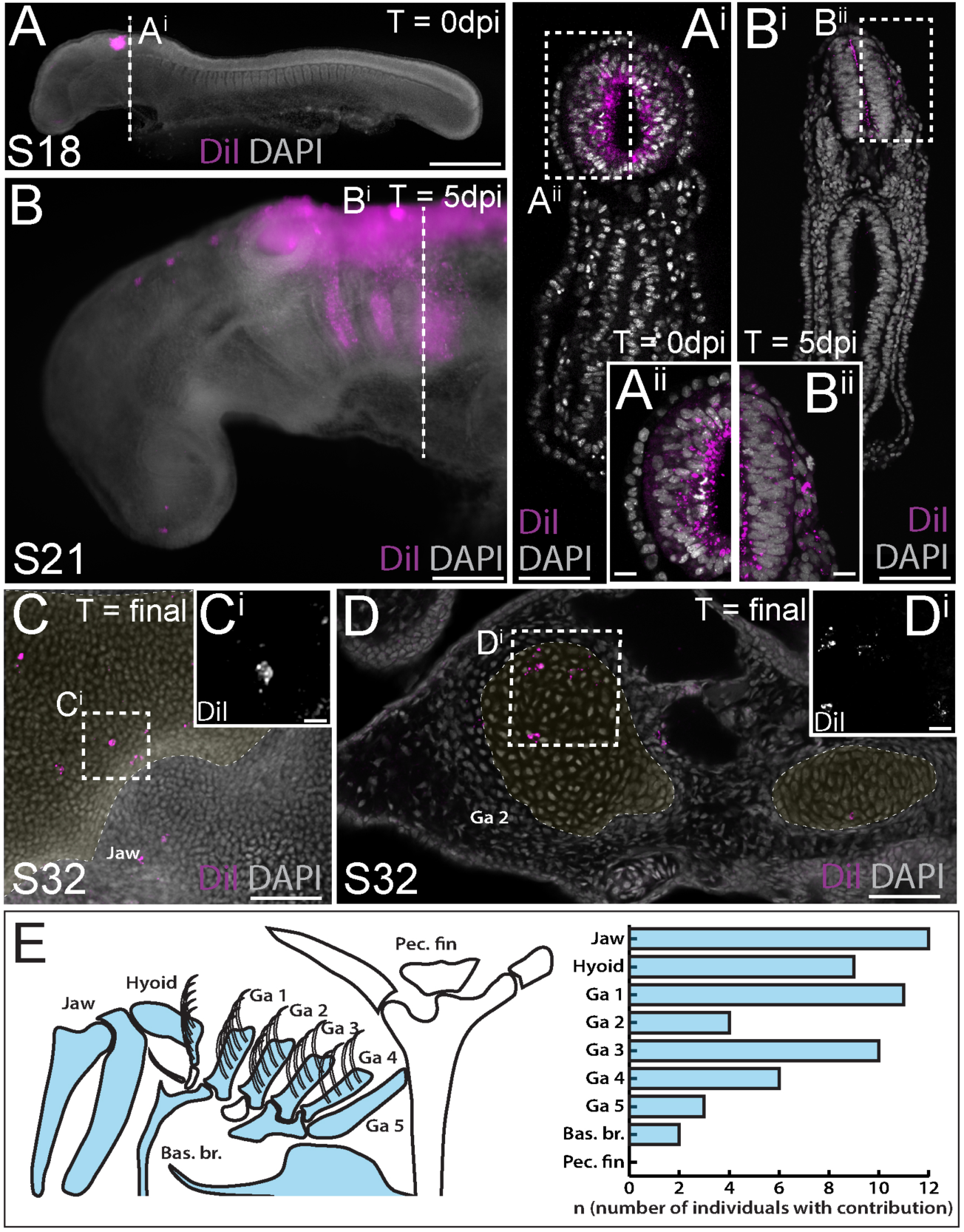
Neural crest contributes to the jaw, hyoid and gill arch skeleton in the skate. **A)** Microinjection of CM-DiI into the lumen of the neural tube at S18 results in **A**^**i**^, **A**^**ii**^) labelling of cells throughout the hindbrain neural tube, including premigratory neural crest cells. **B)** At 5 days post injection (dpi), CM-DiI-labelled cranial neural crest cells can be seen streaming from the hindbrain neural tube into the pharyngeal arches (see also **B**^**i**^, **B**^**ii**^). At S32, CM-DiI (i.e. neural crest-derived) chondrocytes are recovered within pharyngeal arch skeletal elements, including **C, C**^**i**^) the palatoquadrate of the jaw and **D, D**^**i**^) the epibranchial of gill arch 2. **E)** Schematic representation of pharyngeal and pectoral fin skeletal elements in the S32 skate embryo, with elements receiving contribution from neural crest colored blue, and a plot showing the number of embryos observed with neural crest contributions to the pharyngeal arch skeleton. In **C** and **D**, cartilaginous elements are false-colored yellow. Scale bars: **A** = 700μm; **A**^**i**^ = 250μm; **A**^**ii**^ = 50μm; **B** = 340μm; **B**^**i**^ = 250μm; **B**^**ii**^ = 50μm; **C** = 165μm; **C**^**i**^ = 15μm; **D** = 70μm; **D**^**i**^ = 20μm.

We have previously shown that the cartilaginous skeletal elements of embryonic skates can be readily identified, morphologically, in DAPI-stained vibratome or paraffin sections, and that labeling of early embryonic progenitors with lipophilic dyes is an effective way of mapping contributions to the cartilaginous endoskeleton (Gillis *et al*., 2013; Gillis and Hall, 2016; Gillis *et al*., 2017; Criswell et al., 2017; Criswell and Gillis, 2020). While the extent of CM-DiI-labeling of skeletal derivatives is always greatly reduced, relative to the labeling of progenitor cells at the time of injection (due to dilution of the CM-DiI label over several weeks of growth), positively-labeled cells are nevertheless unequivocally recognizable within the skeleton, due to the persistent brightness of the label. To add an additional level of stringency to our analysis, we only scored contributions to the skeleton consisting of clusters of two or more labelled cells, and contributions that were located in the center of a skeletal element (to avoid inadvertently scoring CM-DiI-labeled connective tissue abutting the cartilage). As embryonic cartilage is a homogeneous tissue, consisting of a single cell type (the chondrocyte), we can therefore trace, with great certainty, the contributions of labeled progenitors to the differentiated cartilaginous endoskeleton.

Using the approach outlined above, we readily observed clusters of NC-derived chondrocytes, for example, in the cartilage of the palatoquadrate (Fig. 2C) and the epibranchial and branchial ray cartilages of the first gill arch (Fig. 2D). Overall, our analysis recovered NC contributions to major paired elements of the pharyngeal skeleton (i.e. jaw, hyoid and gill arch elements) and/or to the ventral midline cartilages across all labelled embryos (n=20/20), but no contributions to the pectoral girdle (Fig. 2E; Table S1). These findings are consistent with previous assessments of NC contribution to the pharyngeal and paired fin skeleton of zebrafish using genetic lineage tracing (Kague *et al*., 2012).

### Lateral mesoderm contributes to the skate gill arch and pectoral fin skeleton

We next sought to complement our NC fate map with a test for mesodermal contributions to the pharyngeal and paired fin skeleton in the skate. To do this, we used sub-ectodermal microinjection of lipophilic dyes (CM-DiI or SpDiOC_18_) to label lateral mesoderm at three positions – within the *tbx1*-expressing head mesoderm (HM), at the boundary between HM and *pitx2*/*hand2*-expressing lateral plate mesoderm (LPM), or exclusively within LPM (Fig. 1D) – either alone (Fig. 3A), or in combination with neural crest labelling (Fig. 3B). We once again left labelled embryos to develop for ∼8-10 weeks post-injection, and then scored the embryos for contributions to the skeleton, as described above. Embryos labelled within the HM at S18 showed contributions to pharyngeal arch musculature (Fig. S2), but little contribution to the craniofacial skeleton (labelled chondrocytes were recovered in gill arch cartilage of n=1/10 labelled embryos; Table S1), while in the collective majority of embryos labelled at the HM-LPM boundary (n=10/21) or within the LPM (n=14/17), we observed substantial contributions to the pectoral girdle and fin skeleton (Fig. 3C; Table S1). Remarkably, though, in many embryos labelled at the HM-LPM boundary (n=11/21) or within LPM (n=8/17), we also recovered label-retaining chondrocytes in the skeleton of gill arches 1-5. Mesodermally-derived chondrocytes were recovered within the epi-or ceratobranchial cartilages (e.g. Fig. 3D) and branchial rays (Fig. 3E) of gill arches 1-4, as well as in the ceratobranchial of gill arch 5, in close proximity to the label-retaining pectoral girdle (Fig. 3F – also, see Fig. S3 for additional examples of mesoderm-derived label-retaining chondrocytes within the gill arch skeleton). Overall, our analysis recovered no mesodermal contributions to the mandibular or hyoid arch skeleton, but substantial mesodermal contributions to the paired cartilages of gill arches 1-5, as well as to the pectoral girdle and fin skeleton (Fig. 3G; Table S1).

**Figure 3:**
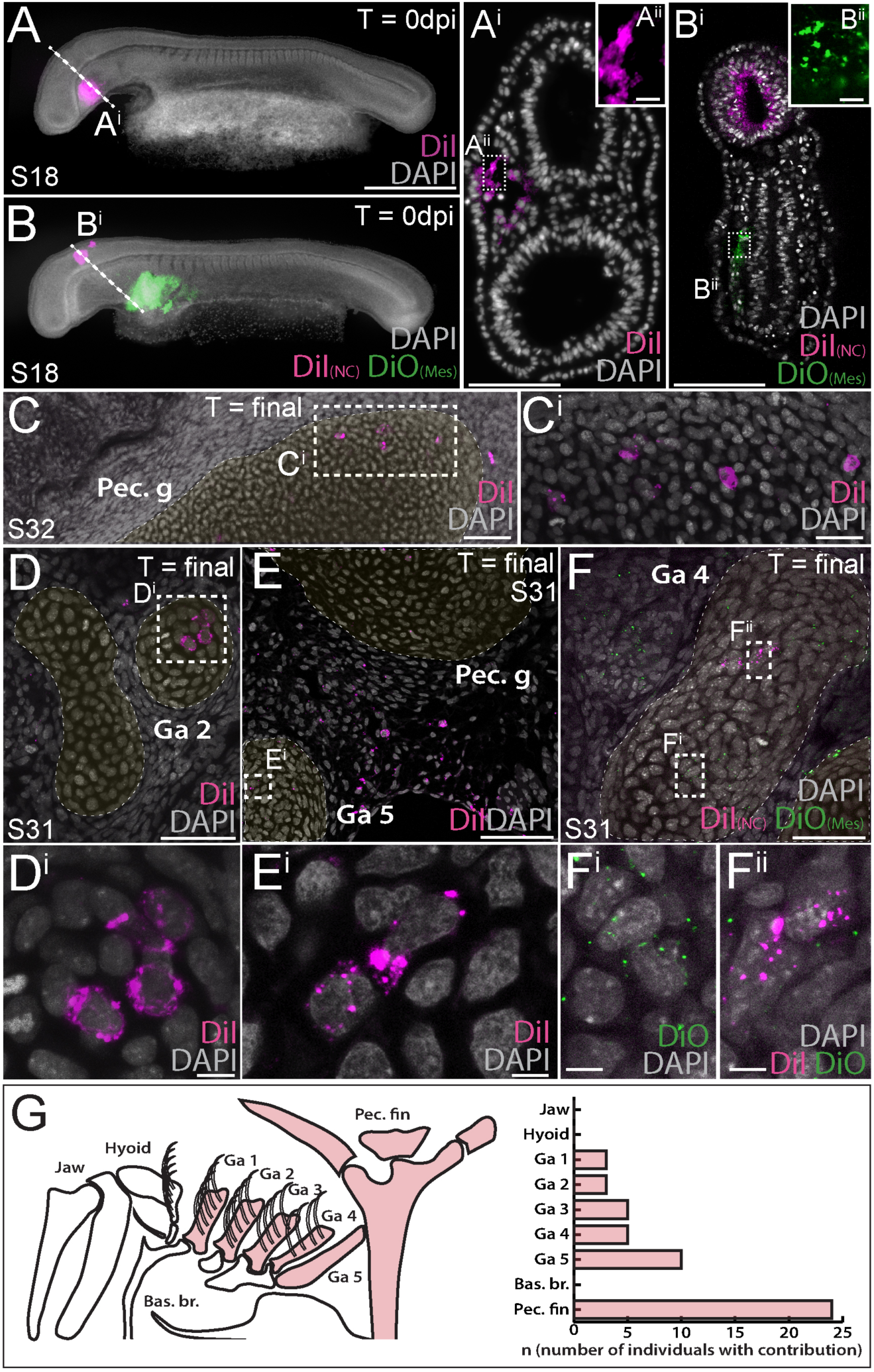
Mesoderm contributes to the gill arch and pectoral fin skeleton in the skate. **A, A**^**i**^) Microinjection of CM-DiI into the head mesoderm (HM) of a skate embryo at S18. **B, B**^**i**^) Simultaneous labelling of the hindbrain neural tube (including premigratory cranial neural crest cells) with CM-DiI and lateral plate mesoderm (LPM) with SpDiOC_18_ in a S18 skate embryo. **C, C**^**i**^) LPM gives rise to chondrocytes within the skeleton of the pectoral fin and girdle, while mesoderm at the HM-LPM boundary and LPM give rise to chondrocytes within the gill arch skeleton – e.g. **D, D**^**i**^) in the ceratobranchial of gill arch 2 and **E, E**^**i**^) the ceratobranchial of gill arch 5. **F)** After double labelling of the LPM with SpDiOC_18_ and the neural tube with CM-DiI, as in **B** above, both **F**^**i**^) SpDiOC_18_- and **F**^**ii**^) CM-DiI-labelled chondrocytes are recovered within the gill arch skeleton – for example, in the ceratobranchial of gill arch 4 –demonstrating the dual mesodermal and neural crest origin of these elements. **G)** Schematic summary of pharyngeal and paired fin skeletal elements in the S32 skate embryo, with elements receiving contributions from mesoderm colored red, and a plot showing the number of embryos observed with mesoderm contributions to the pharyngeal arch and pectoral fin skeleton. In **D, E** and **F**, cartilaginous elements are false-colored yellow. Scale bars: **A, B** = 700μm; **A**^**i**^ = 125μm; **A**^**ii**^ = 15μm; **B**^**i**^ = 50μm; **B**^**ii**^ = 15μm; **C** = 60μm; **C**^**i**^ = 20μm; **D** = 50μm; **D**^**i**^ = 5μm; **E** = 30μm; **E**^**i**^ = 5μm; **F** = 60μm; **F**^**i**^ = 7μm.

## Discussion

When considered alongside lineage tracing data from bony fishes, our findings allow us to infer an ancestral mesodermal contribution to the jawed vertebrate gill arch skeleton (Fig. 4A), with the transition from neural crest-derived to mesodermally-derive skeletogenic mesenchyme occurring gradually, and spanning the region of the posterior (i.e. ancestrally gill-bearing) pharyngeal arches (Fig. 4B). Taken together, our fate mapping experiments point to a neural crest origin of the mandibular and hyoid arch skeleton, a dual NC/mesodermal origin of the gill arch skeleton and an exclusively mesodermal origin of the pectoral fin skeleton in cartilaginous fishes (Fig. 4C). In light of the dual embryonic origin of the mammalian thyroid cartilage and exclusively mesodermal origin of the cricoid and arytenoid cartilages (which are regarded as derivatives of the 4^th^ and 6^th^ pharyngeal arches), it is likely that boundaries of neural crest- and mesodermally-derived skeletogenic mesenchyme have shifted through vertebrate evolution.

**Figure 4:**
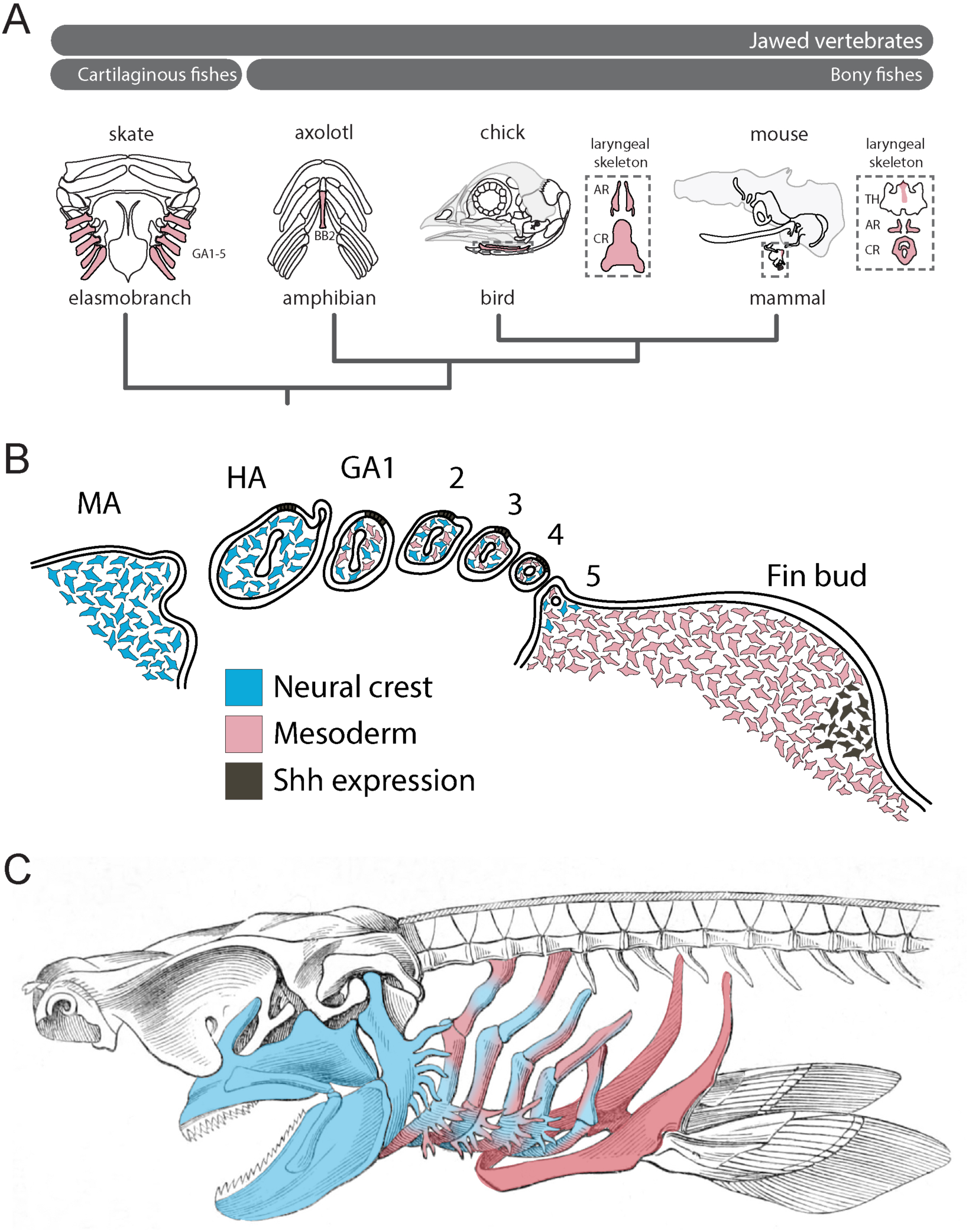
Mesodermal contributions to the pharyngeal endoskeleton in jawed vertebrates. **A)** Mesodermal contributions (red) to the gill arch skeleton in skate, the basibranchial skeleton in axolotl and the laryngeal skeleton of chick and mouse points to an ancestral mesodermal contribution to the pharyngeal arch skeleton of jawed vertebrates. **B)** Schematic representation of neural crest-(blue) and mesoderm-derived (red) skeletogenic mesenchyme in the skate pharyngeal arches and pectoral fin bud, in relation to epithelial and mesenchymal *Shh* expression, respectively. **C)** We propose that the mandibular and hyoid arch skeleton are neural crest-derived and the pectoral fin skeleton mesodermal derived, while the gill arch skeletal elements are of dual neural crest and mesodermal origin.

Our findings also have important implications for understanding the evolutionary origin of paired appendages. With waning support for Gegenbaur’s gill arch hypothesis, the lateral fin fold hypothesis of Balfour (1881), Thatcher (1877) and Mivart (1879) emerged as the favored scenario of paired fin origins. This hypothesis purports that paired fins originated from a continuous epithelial fold that flanked the trunk of the embryo, and that was subsequently segmented into distinct appendages at the pectoral and pelvic levels (reminiscent of the origin of the 1^st^ and 2^nd^ dorsal fins from a continuous median fin fold in sharks). While palaeontological and embryological evidence for the existence of a lateral fin fold (in phylogeny or ontogeny) remains scant, there is evidence of shared molecular patterning mechanisms between dorsal median fins and paired appendages (Freitas *et al*., 2006; Dahn *et al*., 2007; Letelier *et al*., 2018), and of the existence of broad zones of competence along the length of the trunk, from which ectopic fin/limbs or buds may be induced to form (Cohn *et al*., 1995; Kawakami *et al*., 2001; Yonei-Tamura *et al*., 2008). From these observations, a scenario has emerged in which an established appendage patterning developmental module was co-opted, bilaterally, from the dorsal midline to the flank, giving rise to paired pectoral and pelvic appendages.

We previously discovered shared, biphasic roles for Shh signaling in anteroposterior axis establishment and proliferative expansion of skeletal progenitors in the skate hyoid and gill arches and the tetrapod limb bud (Gillis *et al*., 2009; Gillis and Hall, 2016), and we now show that this is regardless of the germ layer origin of Shh-responsive skeletogenic mesenchyme (i.e. neural crest alone in the hyoid arch, neural crest and lateral mesoderm in the gill arches and lateral mesoderm alone in the fin/limb bud) (Fig. 4B). We propose that shared responses of hyoid, gill arch and limb skeletal elements to perturbations in Shh signaling – despite differences in the source of Shh in these organs (i.e. the gill arch epithelial ridge and limb bud zone of polarizing activity – Riddle et al., 1993; Gillis and Hall, 2016) (Fig. 4B) – reflect a common underlying competence of gill arch and fin/limb skeletogenic mesenchyme to respond to these patterning signals, and serial homology of the skeletal derivatives of this mesenchyme. The zones of competence that underlie the origin of pectoral and pelvic appendages within the trunk could, accordingly, be extended rostrally to include zones of neural crest and mixed neural crest/lateral mesodermal contribution to the pharyngeal endoskeleton, and this, in turn, could account for the serial derivation of gill arches and paired appendages along the gnathostome stem. Indeed, reports of a fossil jawless vertebrate with gill arches extending down the length of the trunk (Janvier *et al*., 2006) further support the shared competence of pharyngeal and lateral trunk mesenchyme to give rise to both gill arch and fin/limb skeletal elements.

Importantly, a competence-based hypothesis of gill arch-fin serial homology decouples the origin and evolutionary histories of gill arches/paired appendages as anatomical structures and the molecular mechanism that direct their patterning – i.e. it accounts for the former, but leaves the latter open to further discourse around the deep homology of appendage patterning mechanisms within vertebrates or, more broadly, metazoans (Shubin *et al*., 2009). It is widely appreciated that, in animals, a relatively small number of developmental signaling pathways are used repeatedly, and in different combinations/contexts, to instruct the development of a great many embryonic tissues and organs. This, in turn, precludes the straightforward inference of homology of anatomical structures based on shared molecular patterning mechanisms (Dickinson, 1995). We argue that recognition of anatomical similarity due to common *response* to instructive cues within generative tissues, rather than focusing on the cues themselves, can allow us to bridge the gap between patterning mechanisms and morphology, and may provide a basis for inferring homology of morphology, even when considering structures that develop under the influence of upstream patterning mechanisms with complex and/or distinct evolutionary histories.

Homology is a hierarchical concept, and two complex features (e.g. organs) – which arise within the context of an embryonic tissue, by deployment of a gene regulatory network operating downstream of an inductive or patterning cue – may be homologous at one biological level of organization, while simultaneously non-homologous at another (Hall, 2003; Wagner, 2014). While reconstructing the evolutionary history (homology) of individual genes or gene regulatory network nodes is becoming increasingly straightforward, meaningfully testing the homology of putatively distantly-related structures at the anatomical level – whether historical homologues across taxa, or serial homologues within a taxon – has, in many cases, lingered as problematic. Developmental competence, or the cell-autonomous property that imparts on tissues an ability to respond to external stimuli (e.g. organizers) (Waddington, 1947), may represent a tangible means of linking upstream molecular developmental mechanisms with ultimate anatomical readouts (Spemann, 1915). In light of the demonstrated lability of germ layer fates within the vertebrate skeleton (Schneider, 1999; Teng *et al*., 2019), we suggest that, in the case of anatomy, competence – which is inherently testable, either by natural (i.e. evolutionary) or laboratory experimentation – may supersede germ layer origin as a primary criterion of homology.

## Materials and Methods

### Embryo collection

*L. erinacea* eggs were obtained at the Marine Biological Laboratory (Woods Hole, MA, USA) and maintained in a flow-through seawater system at ∼15°C to the desired developmental stage. Embryos for mRNA *in situ* hybridization were fixed in 4% paraformaldehyde in phosphate-buffered saline (PBS) overnight at 4°C, rinsed three times in PBS, dehydrated into 100% methanol and stored at −20°C. Embryos injected with CM-DiI and SpDiOC_18_ were fixed in 4% paraformaldehyde in PBS overnight at 4°C, rinsed three times in PBS, and stored in PBS + 0.02% sodium azide at 4°C.

### mRNA in situ hybridization

*L. erinacea* embryos were embedded in paraffin wax and sectioned at 8 μm thickness for mRNA *in situ* hybridization as previously described (O’Neill *et al*., 2007). Wholemount and paraffin chromogenic mRNA *in situ* hybridization experiments for *FoxD3* (GenBank accession number MN478366), *Tbx1* (GenBank accession number MT150581), *Pitx2* (GenBank accession number MT150579), *Hand2* (GenBank accession number MT150580) and *Myf5* (GenBank accession number MT150582) were performed as previously described (O’Neill *et al*., 2007) with modifications according to Gillis *et al*. (2012).

### Fate mapping and imaging

Preparation and microinjection of CM-DiI and SpDiOC_18_ was carried out as previously described (Gillis *et al*., 2017; Criswell and Gillis, 2020). After labelling, sealed eggs were returned to a flow-through seawater system at ∼15°C to the desired developmental stage, and then euthanized using an overdose of tricaine (1g/L in seawater) prior to fixation. Labelled embryos to be analyzed by vibratome sectioning were rinsed 3×5 minutes in PBS, embedded in 15% (w/v) gelatin in PBS and post-fixed in 4% paraformaldehyde in PBS for 4 nights at 4°C before sectioning at 100μm on a Leica VT1000S vibratome. Sections were then DAPI-stained (1μg/mL), coverslipped with Fluoromount-G (Southern Biotech) and imaged on an Olympus FV3000 confocal microscope. Labelled embryos to be analyzed by paraffin histology were embedded and sectioned as previously described (O’Neill *et al*., 2007).

## Acknowledgments

The authors thank Dr. Richard Schneider, Prof. David Sherwood, and the MBL Embryology Course for provision of lab space, Louise Bertrand and Leica Microsystems for microscopy support, the staff of the Marine Resources Center at the MBL for assistance with animal husbandry and Dr. Kate Criswell, Jenaid Rees, Christine Hirschberger and Dr. Kate Rawlinson for helpful discussion. This project benefited from technical advice from Dr. Matt Wayland and use of the Imaging Facility, Department of Zoology, supported by a Sir Isaac Newton Trust Research Grant (18.07ii(c)). This research was supported by a Royal Society University Research Fellowship (UF130182) and grants from the Leverhulme Trust (RPG-2016-373) and the University of Cambridge Sir Isaac Newton Trust (14.23z) to JAG, and by a Junior Research Fellowship from Wolfson College, Cambridge and Whitman Early Career Fellowship from the Marine Biology Laboratory to VAS.

## Author contributions

JAG conceived and oversaw the study. VAS contributed to experimental design, performed, analyzed and imaged all experiments, and prepared all figures. VAS and JAG interpreted the data. JAG wrote the manuscript with input from VAS.

---

**Figure S1:**
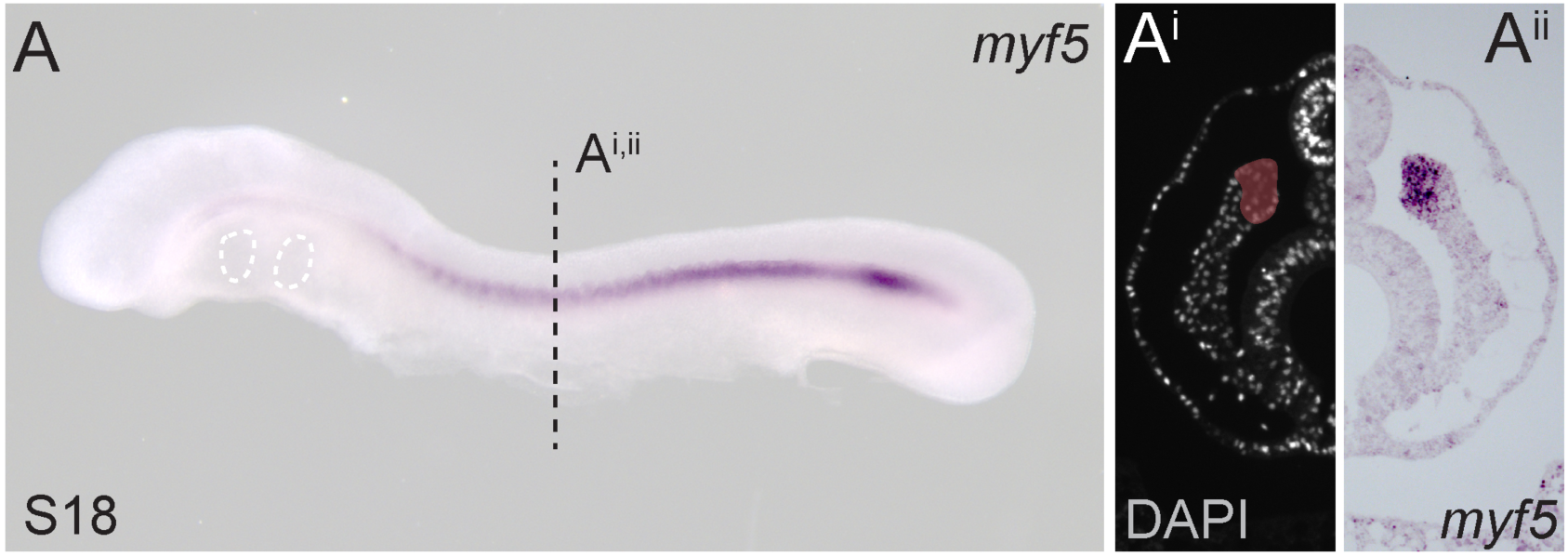
Paraxial mesoderm in the little skate. **A)** Wholemount mRNA *in situ* hybridization for *myf5* reveals expression in **A**^**i**^, **A**^**ii**^) somitic and presomitic mesoderm of the skate embryo at S18.

**Figure S2:**
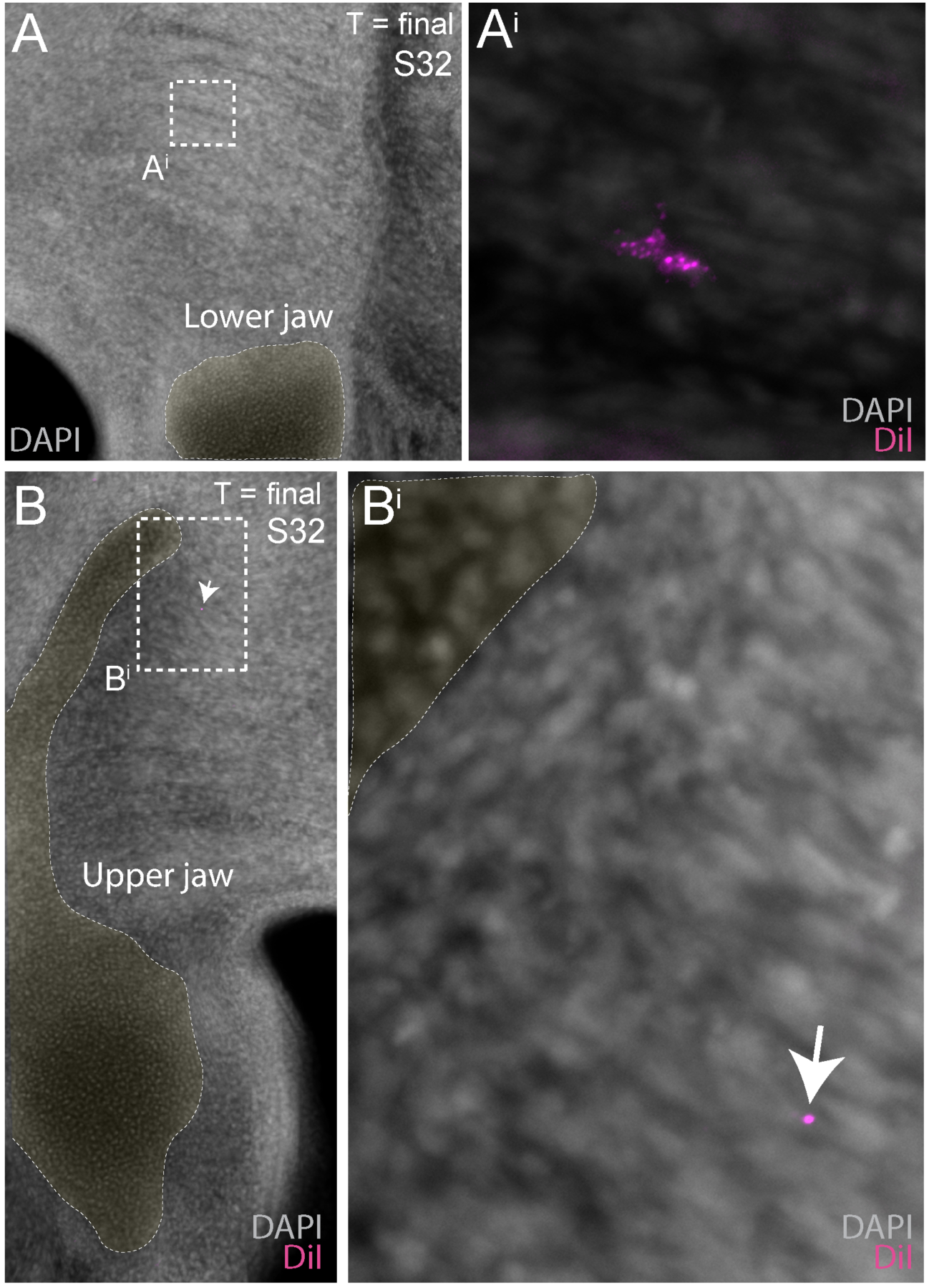
Head mesoderm contributes to pharyngeal arch musculature in the skate. In skate embryos in which head mesoderm was labelled with CM-DiI at S18, CM-DiI-labelled cells were subsequently recovered in **A, A**^**i**^) lower jaw and **B, B**^**i**^) upper jaw musculature. Lower and upper jaw cartilages are false-colored yellow.

**Figure S3:**
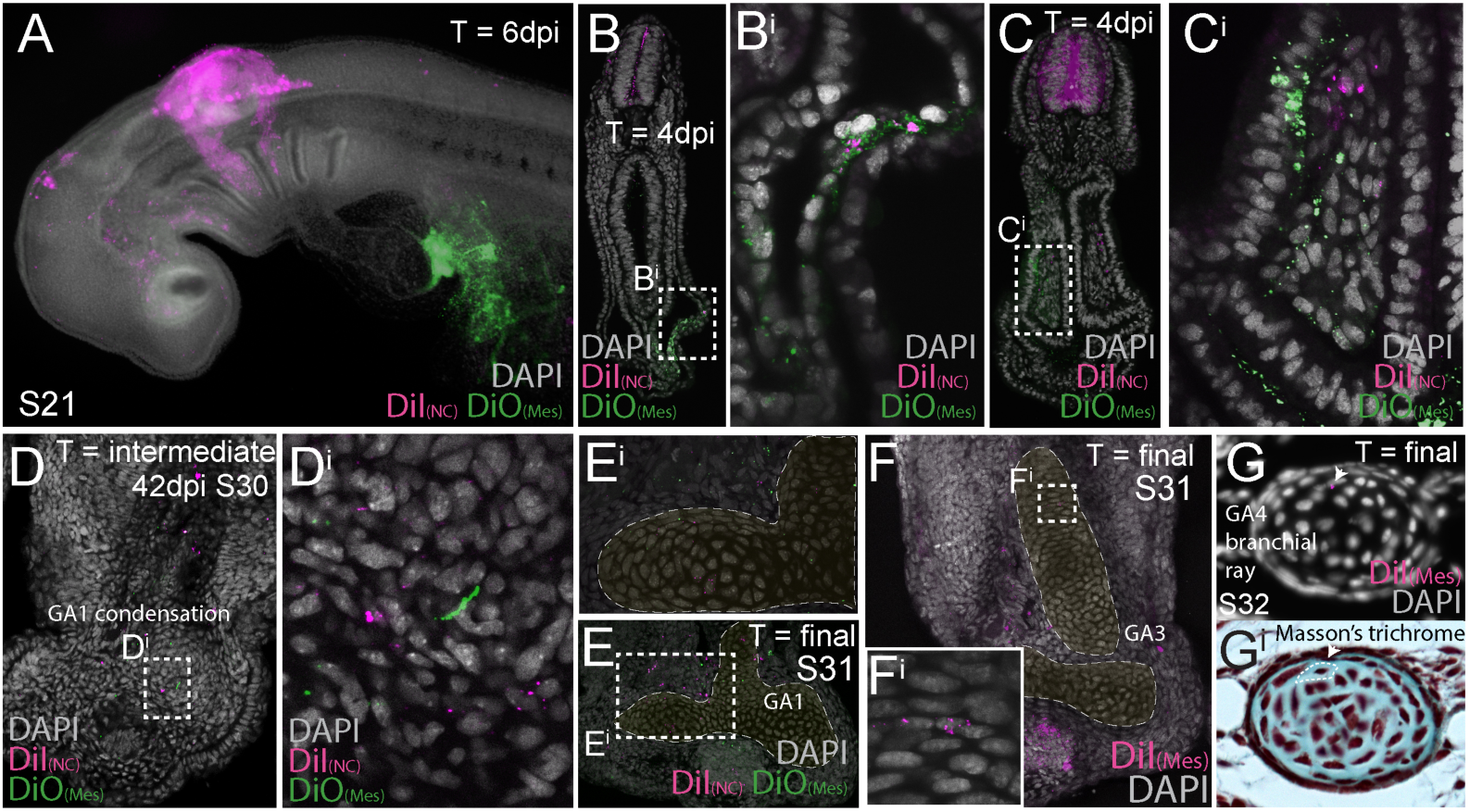
Dual embryonic neural crest and mesodermal origin of gill arch cartilages in the skate. **A)** A skate embryos 6 days after neural tube labelling with CM-DiI and lateral plate mesoderm (LPM) labelling with SpDiOC_18_. **B-C)** Transverse sections reveal that 4 days post neural tube labelling with CM-DiI and LPM labelling with SpDiOC_18_, CM-DiI-labelled neural crest cells can be seen migrating adjacent to SpDiOC_18_-labeled LPM. **D)** Forty-two days post dual labelling, both CM-DiI-retaining (i.e. neural crest-derived) and SpDiOC_18_-retaining (i.e. LPM-derived) cells are recovered within the condensing ceratobranchial cartilage of gill arch 1, while **E)** at S32, CM-DiI-retaining and SpDiOC_18_-retaining chondrocytes are recovered within the ceratobranchial cartilage of gill arch 1. **F)** Mesodermal contribution to the ceratobranchial cartilage of gill arch 3. **G)** Mesodermal contribution to a branchial ray on gill arch 4. In **E** and **F**, cartilaginous elements are false-colored yellow.

**Data S1. (separate file)**

Complete scoring table for neural crest and mesodermal fate mapping experiments

